# Role of oligodendroglial ADAM10 in oligodendrocyte maturation, myelination and myelin maintenance

**DOI:** 10.1101/2025.07.25.666787

**Authors:** Mathis Lavaud, Aïda Padilla-Ferrer, Anne Simon, Pilar Acebo, Celine Dargenet-Becker, Victor Gorgievski, Eleni Tzavara, Charbel Massaad, Mehrnaz Jafarian-Tehrani, Delphine Meffre

## Abstract

In the central nervous system, oligodendrocytes extend their processes to wrap axons to form myelin sheaths, ensuring saltatory conduction of action potentials and providing trophic support for the axon. Our previous work demonstrated that pharmacological activation of α-secretase promotes myelin protection and remyelination after both *ex vivo* and *in vivo* demyelination. Based on these findings, the present study aims to investigate the role of oligodendroglial ADAM10 (OLA10; a member of the a-secretase family) in myelin development and maintenance. Using an inducible knockout mouse model for OLA10 (KOOL-A10), we demonstrated that OLA10 deficiency results in delayed maturation of OPC in primary culture, and altered myelination in primary neuron/glia co-culture. Furthermore, an induction of OLA10 deficiency in adult mice was associated with an alteration of myelin sheath thickness and long-term motor and cognitive deficit in males but not in females. Overall, our study highlights the subtle but significant role of ADAM10 expressed by oligodendrocytes in the formation and maintenance of the myelin over time, while emphasizing a sexual dimorphism that carries important relevance for current research.

## 1- Introduction

Multiple sclerosis (MS) is a neurodegenerative demyelinating disease that targets the central nervous system (CNS) (Dobson & Giovannoni, 2019). One of the main hallmarks of MS is demyelination, a process characterized by the loss of the myelin sheath and/or oligodendrocyte, leading to impaired saltatory conduction and subsequent progressive axonal degeneration. Following demyelination, spontaneous remyelination occurs in an attempt to restore the integrity of the myelin sheath and protect axons. In the context of MS, remyelination is disrupted and results in axonal damage (Schäffner et al., 2023). Several studies suggest that oligodendrocyte differentiation is a critical step for the remyelination process (Dulamea, 2017; Kornberg & Calabresi, 2024), making the development and maturation of oligodendrocyte progenitor cells (OPCs) one major research focus on spontaneous remyelination in MS (Dulamea, 2017; Ogata, 2019; Tepavčević & Lubetzki, 2022).

We have previously demonstrated the beneficial effects of etazolate treatment both *in vivo* and *ex vivo* in models of cuprizone-, trauma- or lysolecithin-induced demyelination (Llufriu-Dabén et al., 2018, 2019; Carrete et al., 2021). Our data revealed the protective effects of this pharmacological compound, as well as its promyelinating action, along with a differentiation-inducing effect on oligodendrocytes. Etazolate is known to act as an activator of α-secretases, an enzyme family including different members of ADAMs (A Disintegrin And Metalloproteinase), such as ADAM9, 10 and 17. ADAMs are transmembrane proteases synthesized as precursors and matured into their active forms with multiple substrates, exerting important biological roles via their effects on cell adhesion, migration, proteolysis and signaling (Edwards et al., 2008). Several studies showed that these enzymes could be involved in myelination and myelin maintenance (Hsia et al., 2019). For example, in the peripheral nervous system (PNS), the neuronal α-secretase ADAM17 has been shown to be involved in PNS myelination (La Marca et al., 2011), while the ADAM17 expressed in Schwann cell contributes to remyelination (Pellegatta et al., 2022). In the CNS, the neuronal ADAM17 has been reported to play a role in OPC development (Fredrickx et al., 2020), and the glial form in both myelination and remyelination (Palazuelos *et al*., 2014, 2015). ADAM17 shares some of its substrates with another α-secretase, ADAM10. In the PNS, ADAM10 does not appear to play a direct role in myelination but does affect axonal growth (Jangouk et al., 2009; Luo et al., 2011; Meyer Zu Horste et al., 2015). However, some findings suggested that its neuronal form could influence nerve myelination *in vitro* through one of its substrates, DR6 (Colombo et al., 2018). ADAM10 has been characterized and mapped within the CNS, where it is expressed in various cell types including oligodendrocytes (Guo et al., 2016; Padilla-Ferrer et al., 2024). Neuronal ADAM10 has been extensively studied in relation to its various substrates in the CNS (Lin *et al*., 2008), and total knockout (KO) of ADAM10 is lethal at embryonic stage, highlighting important CNS alterations (Hartmann *et al*., 2002). ADAM10 also exhibits neuroprotective properties via its enzymatic activity, notably by cleaving APP (Amyloid Precursor Protein) into sAPPα (soluble Amyloid Precursor Protein alpha) (Mockett *et al*., 2017; Vitória *et al*., 2022). In post-mortem CNS tissues from MS patients, ADAM10 was predominantly detected in astrocytes and in perivascular macrophages within chronic active plaques (Kieseier *et al*., 2003). While these studies did not primarily address myelination or remyelination, Zhu and collaborators showed that ADAM10 overexpression with no specific cell targeting in the corpus callosum via adenoviral delivery exerts neuroprotective and remyelinating effects comparable to sAPPα overexpression following cuprizone-induced demyelination (Zhu *et al*., 2021). The neuronal ADAM10 activity may explain, at least in part, the beneficial effect since it is required for the migration of neural precursor cells into demyelinated lesions (Klingener *et al*., 2014). Another group using a NG2 glial-specific KO of ADAM10, targeting both astrocyte and oligodendrocyte, reported an earlier onset of myelination accompanied by behavioral alterations (Guo *et al*., 2022). It has been also reported that ADAM10 expression in glial progenitors regulates oligodendrogenesis and astrogenesis in the developing brain (Wang *et al*., 2023). ADAM10 thus appears to be a promising target in the processes of myelination and remyelination. Having previously identified ADAM10 expression in oligodendrocytes (OLA10) (Padilla-Ferrer *et al*., 2024), we aim to explore the role of OLA10 in myelination and myelin maintenance.

To investigate the effect of OLA10 deficiency on oligodendrocyte maturation and myelin maintenance, we developed a conditional and inducible KO mouse model for OLA10, referred to as KOOL-A10. We first assessed the effects of OLA10 deficiency *in vitro* in primary OPC cultures to evaluate OPC morphological maturation, and then in neuron/glia co-cultures to examine the timing of myelination. Subsequently, OLA10 was inactivated in adulthood to enable a longitudinal analysis spanning from 1 to 12 months post-invalidation, aimed at investigating the impact of OLA10 deficiency on myelin maintenance. We analyzed the ultrastructure of the myelin sheath in cerebellum *in vivo* using electron microscopy, and the level of the major myelin protein MBP (Myelin Basic Protein) in brain, cerebellum and spinal cord. Finally, we evaluated whether the absence of OLA10 led to alterations in motor and cognitive functions with appropriate behavioral tests. Our results show that OLA10 deficiency disrupts the morphological maturation of oligodendrocytes and their ability to form myelin *in vitro*. We also observed changes in the myelin ultrastructure *in vivo*, accompanied by impairments in motor and cognitive behavior, with sexual dimorphism.

## 2- Experimental procedures

### 2.1- Animals

Animal care and experiments were approved by the Université Paris Cité Animal Ethics Committee (APAFIS #16685, #47419 and #47420). They adhere to French regulations as well as the European Communities Council Directive on the protection of animals used for scientific purposes. Mice were housed at 22 ± 2°C with a 12-hour light/dark cycle. KOOL-A10 mouse strain was obtained from the breeding of two strains: ADAM10-flox strain (B6; 129S6-Adam10^tm1Zhu^/J), where exon 3 of *Adam10* gene is flanked by loxP sites (Tian *et al*., 2008), and PLP-creER^T2^ strain (B6.Cg-Tg(Plp1-cre/ERT)3Pop/J) (Doerflinger *et al*., 2003). Mice were used at E13.5 for co-culture (**Fig. 1B**), at P3-P4 for OPC primary culture (**Fig. 1A**) and at adult stage for *in vivo* studies (5-months-old male and female mice to start tamoxifen treatment for the induction of OLA10 deficiency and multiple analysis at 1, 6 or 12-months post-treatment, **Fig. 1C**). Tamoxifen (Sigma, T5648) was diluted in sunflower oil and administered either by pipette feeding once at P0 (100 mg/kg) or by intraperitoneal injection in adults (60 mg/kg) for 3 consecutive days. ADAM10-flox mice (KOOL-A10 littermates without Cre) treated with tamoxifen were used as control mice. For histological studies, mice were perfused using phosphate buffer at 0.1 M (pH 7.4) and then fixated using 4% PFA (Sigma, P6148). Brain was immersed in a cold fixative solution (4% PFA, 2.5% glutaraldehyde and 0.1 M phosphate buffer, pH 7.4) and stored at 4°C until use. For biochemical studies, mice were sacrificed by cervical dislocation, and the tissues were removed, snap-frozen in liquid nitrogen, and stored at −80°C until use.

**Figure 1:**
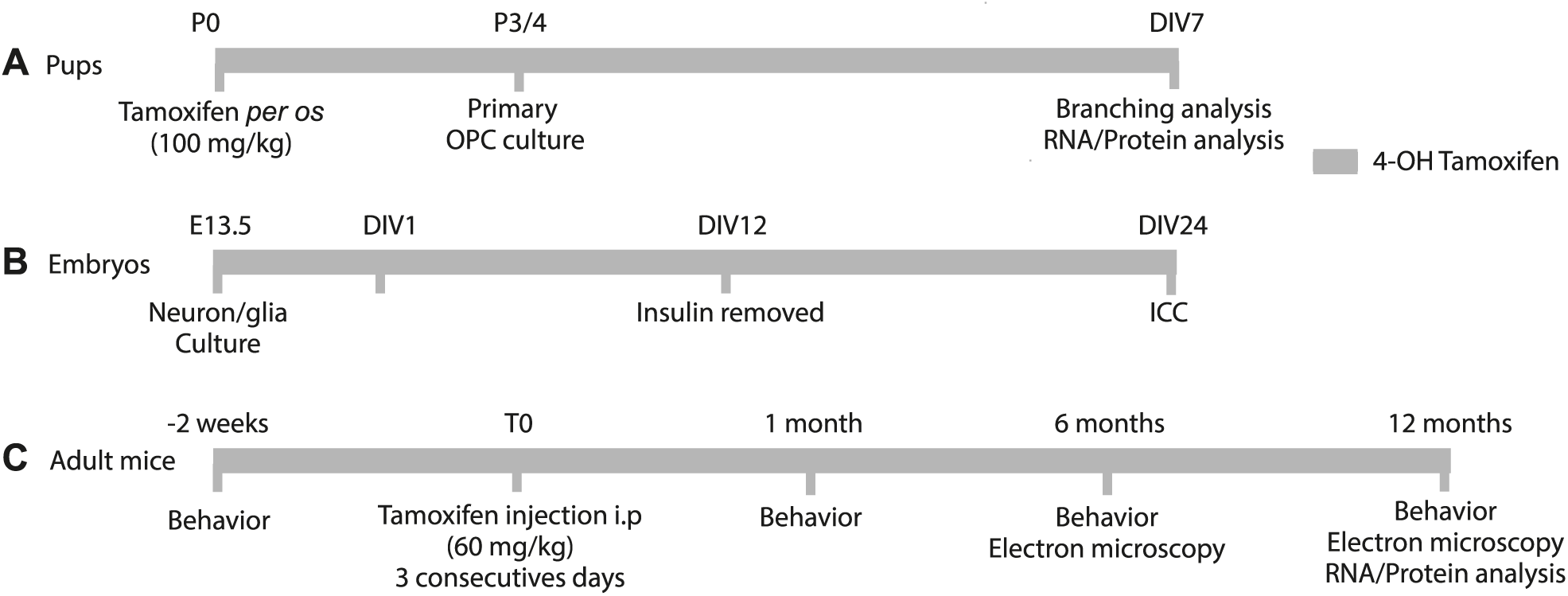
Experimental timelines. **(A) OPC primary culture experimental timeline.** KOOLA-10 pups were treated *per os* with tamoxifen at P0 (100 mg/kg). Brain and cerebellum were removed at P3-P4, dissociated to isolate OPCs using MACS® technology and cells were cultivated with 4-OH-tamoxifen (1 µM). At DIV7, pictures were acquired with inverted brightfield microscope (20X) before RNA/proteins being extracted. **(B) Neuron/glia co-culture experimental timeline.** Cells from E13.5 KOOLA-10 spinal cords were isolated. Co-cultures were treated with 1 µM of 4-OH-tamoxifen at DIV1, and insulin was removed at DIV12 to initiate differentiation. Cells were fixed with 4% PFA at DIV24 to perform immunocytochemistry experiments. **(C) Experimental timeline for *in vivo* studies.** Adult KOOL-A10 mice were injected with tamoxifen (60 mg/kg, i.p.) during 3 consecutive days. Mice were tested for motor and cognitive abilities, prior tamoxifen, and after (1, 6, 12 months). Histological and biochemical analysis were also performed. DIV: Days *in vitro*, E13.5: Embryonic day 13.5, ICC: Immunocytochemistry, i.p: intraperitoneal injection, OPC: Oligodendrocyte Precursor Cells, P: Postnatal day, T: Time.

### 2.2- Cell cultures

#### 2.2.1- Oligodendrocyte primary culture

P3-P4 KOOL-A10 pups were pretreated with tamoxifen at P0 as described above. Brain hemispheres and cerebella were mechanically dissociated using neural dissociating kit™ (NTDK, Miltenyi, 130-092-628) using GentleMACS™ with "37C_NTDK_1" dissociation program. Cells were incubated with FC receptor blocking solution (10 min, 4°C), then with Microbead Kit™ (anti-CD140a, *i.e.* anti-PDGFRα, Miltenyi, 130-101-502, 15 min, 4°C), and applied on magnetic LS columns™ (Miltenyi, 130-122-729) in MultiMACS 24™ using "POSSEL-SCA" program. PDGFRα^+^ cells were eluted with culture media containing MACS Neuromedium™ (Miltenyi, 130-093-570), 2% Neurobrew (Miltenyi, 130-093-566), 1% Penicillin and Streptomycin (ThermoFisher, 15140122), Glutamax 1X (Gibco, 35050038), 1 μM 4-OH-tamoxifen (Enzo, ALX-550-361-M005), PDGF-AA (10 ng/ml, Peprotech, 315-17) and Human FGF (10 ng/ml, Sigma, F0291). Cells were seeded at 125 cells/mm² in 6 wells plates coated with Poly-L-Lysine (25 μg/ml, Sigma, P5899) and cultivated with 1 µM of 4-OH-tamoxifen until seven days *in vitro* (DIV7) at 37°C and 5% CO_2_. For biochemical analysis, cells were collected in TRIzol™ (Invitrogen, 15596-018) or RIPA buffer containing 50 mM Trizma base pH=7.4, 150 mM NaCl, 1mM MgCl_2_, 1% NP-40, 0.1% SDS, 0.5% Na deoxycholate, Protease inhibitor (Merck, 11836170001), 10 mM Pyrophosphate (Sigma, S9515), 1% Phosphatase inhibitor cocktail 2 and 3 (Sigma Aldrich, P5726 and P0044), 10 mM NaF (Sigma, S7920), and 5 μM GI254023X (Tocris Biosciences, 3995). For branching analysis, 3 images/well were taken at DIV7 with an inverted brightfield microscope using 20X magnification (Leica, DMi FOV20). Cells were classified in 2 types based on their branching: low (1-4) and high degree of ramification (>4 processes) and were counted with *cell counter* plug-in on ImageJ software (1.52k).

#### 2.2.2- Primary neuron/glia co-culture

E13.5 KOOL-A10 spinal cords were dissociated using trypsin (Sigma, T4299) and DNAse (0.5 mg/ml, Roche, 11284932001). 125 000 cells/slide were seeded onto poly-L-lysine coated (100 µg/ml, Sigma, P1274) glass slides (Euromedex, EM-72290-05) with plating media containing 50% low glucose DMEM (Gibco, 31885023), 25% horse serum (Gibco, 26050089) and 25% HBSS (Gibco, 140025051) supplemented with Glucose (19.45 mM, Sigma, G8769), Progesterone (20 nM, Sigma, P0130), Selenium (5.2 ng/ml, Sigma, S9133), Putrescine (16.1 µg/ml, Sigma, P7505), Apotransferrine (5 µg/ml, Sigma, T2252), Biotin (10 ng/ml, Sigma, B4501), Hydrocortisone (50 nM, Sigma, H0396), Insulin (10 µg/ml, Sigma, I0516), Penicillin/Streptomycin (1X, Gibco, 15070-063) and Fungizone (0.5 µg/ml, Gibco, 15290026). Cells were treated at DIV1 with 1 µM of 4-OH-tamoxifen (Enzo, ALX-550-361-M005). Insulin was removed at DIV12 to induce differentiation, and they were maintained in culture until DIV24 (37°C, 5% CO_2_).

### 2.3- RNA extraction and RT-qPCR

RNA was extracted from DIV7 OPCs using TRIzol™ (Invitrogen, 15596-018), and from adult tissues by homogenization in ceramic beads tubes (Fisherbrand, 15555799) using a BeadMill24™ agitator (FisherBrand). Following the addition of chloroform at a 1:5 ratio, samples were centrifugated and 100% ethanol was added at 2.5:1 ratio, supplemented with glycogen (20 mg/ml, Sigma, G1767). After overnight incubation at -20°C, samples were centrifugated (16 000 g, 15 min, 4°C) and washed 4 times with 70% ethanol. Pellets were air-dried (RT, 30 min) and resuspended in RNAse/DNAse free water (Invitrogen, 10977-035). Concentration and RNA purity were assessed by UV spectrophotometry with NanoDrop one™ (Thermo Scientific). For reverse transcription, M-MLV-RT enzyme (Sigma, M1302) with random primers (7 ng/µl, Promega, C1181), dNTPs (10 nM, Promega, U1205, U1225, U1215, U1235) and 250 ng of RNA were used. qPCR was performed with Absolute SYBR ROX 2X qPCR™ kit (Thermo Scientific™, AB1162B) and specific primers (**Table 1**). All samples were run in triplicate and each experiment included a no-cDNA control and a no-RT enzyme control. To assess the specificity of the amplification, melt curve analysis was performed. A single peak with a melting temperature (Tm)> 75°C was expected. Cycles quantification (Cq) values were determined by linear regression, and standard deviation (SD) was then computed across triplicates. Samples presenting a SD > 0.30 were excluded. Gene expression was normalized using housekeeping genes as described in (Sundaram *et al*., 2019): *Ppia* and *Gapdh* (brain), *26s* and *Mrlp10* (cerebellum), *Hsp60* and *Mrlp10* (spinal cord); *26s* and *Ppia* (OPCs). Fold change were computed with the comparative methods 2^−(ΔΔCt)^ (Livak & Schmittgen, 2001; Schmittgen & Livak, 2008) and expressed in percentage of induction relatively to the appropriate control.

**Table 1:**
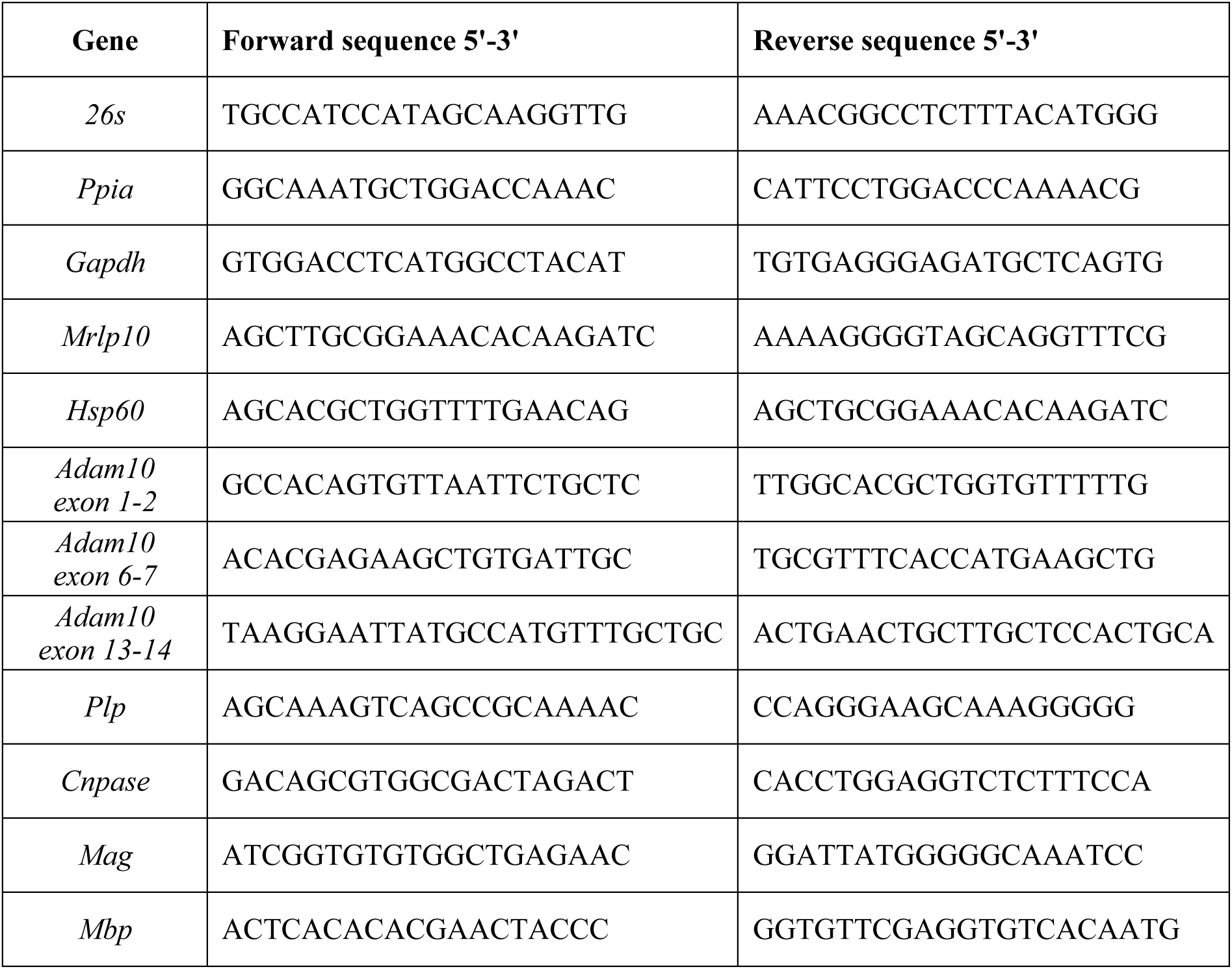
Primers used in RT-qPCR experiments. Sequences are listed as 5’-3’. Primers were used at 300 nM. *26s*: ribosomal RNA, *Adam10*: A Disintegrin and Metalloprotease 10; *Gapdh*: Glyceraldehyde 3-phosphate dehydrogenase; *Cnpase*: 2’,3’-Cyclic-nucleotide 3’ phosphodiesterase; *Hsp60*: Heat shock protein 60; *Mag*: Myelin-associated glycoprotein; *Mrlp10*: Mitochondrial ribosomal protein L10; *Plp*: Proteolipid Protein; *Ppia*: Peptidylprolyl isomerase A.

### 2.4- Protein analysis

Samples were homogenized in RIPA buffer (cf. 2.2.1 for composition) and centrifuged (15 000g, 20 min, 4°C). Protein concentrations were determined using the DC protein assay kit (Biorad, 5000116) with BSA (Sigma, A7906) as the standard reference. After denaturation (95°C, 5 min), 10 µg (cells) or 50 µg (tissues) of protein were separated on a polyacrylamide gel (4-20% Precast gel Biorad, 4561024) by electrophoresis and transferred on an ethanol activated PVDF (Polyvinylidene fluoride) membrane (Biorad, 10026933). Transfer was performed with Trans-Blot® Turbo RTA Midi 0.2 µm PVDF Transfer Kit (Biorad, 1704273) and Trans-Blot turbo transfer system (Biorad, 1704150) using the "High molecular weight" transfer program. After blocking (TBS/5% BSA), membranes were probed overnight at 4°C with the following primary antibodies: Actin (1:10 000, Abcam ab8227), MBP (1:500, Millipore MAB381) and then with the appropriate horseradish peroxidase-conjugated secondary antibodies: goat anti-rabbit (Jackson IR 111-035-144) or goat anti-mouse (Jackson IR 115-035-003), both at 1:20 000 (2h, RT), followed by ECL detection (Amersham, RPN2209). Images were acquired using the ImageQuant™ Las4000 (GE Healthcare) and relative protein amounts were quantified using ImageJ (Gel Analyzer plugin) and normalized to Actin. When necessary, membranes were stripped for 15-20 min at RT using stripping buffer (Thermo Scientific, 21059) prior to reprobing.

### 2.5- Cytology and histology

#### 2.5.1- Immunocyto/histochemistry and imaging

Brains from perfusion-fixated mice were dissected and sectioned coronally in 20 μm thickness using a frozen microtome (Leica SM2010R). Sections (ranging from -1.46 mm to -1.7 mm from Bregma) were incubated in a blocking solution (PBS-GTA) containing PBS (1X), 0.1 M lysine (Sigma, L5628), 0.25% gelatin, 0.2% Triton 100X (Sigma, T8787) and 0.1% Azide. Sections were then incubated overnight at 4°C with the following primary antibodies diluted in PBS-GTA: ADAM10 (1:2 500 (Proteintech, 25900-1), Olig2 (1:1 000, R&D, AF2418), NeuN (1:1 000, Merck MAB377), MAG (1:1 000, Merck, MAB1567), CaBP (1:1 000, Swant, CD38a) and NFH (1:1 000, Merck, AB1989). Then, the appropriate secondary antibodies purchased from Thermo Fisher were used at 1:1 000, for 2h at RT: donkey anti-mouse Alexa Fluor™ Plus 488 (A32766), donkey anti-rabbit Alexa Fluor™568 (A10042), donkey anti-rabbit Alexa Fluor™ Plus 647 (A32795) and donkey anti-goat Alexa Fluor™ Plus 488 (A32814). Sample were mounted with Fluoromount-G™ (Thermo Fischer Scientific, 00-4958-02) with bis- Benzimide™ (1:100 000, Sigma, 14530). For the co-cultures, fixed at DIV24 with 4% PFA, the same protocol was used, except that blocking solution was supplemented with 3% donkey serum. All confocal images were acquired using a Zeiss LSM710 confocal microscope with a 20X Plan-Apochromat/0.8 M27 objective. To determine the number of ADAM10^+^ cells in the brain, thalamic area was selected in order to better discriminated oligodendrocytes from neurons. The number of total NeuN^+^ and Olig2^+^ cells, as well as the co-stained ADAM10^+^ NeuN^+^ and ADAM10^+^ Olig2^+^cells were counted using ImageJ software (1.52k). To assess myelination in co-cultures, a line was drawn across each image, positioned to intersect the highest possible number of axons, using the same software. Along this line, both the total number of axons and the number of myelinated axons were counted. The percentage of myelinated axons was then calculated by dividing the number of myelinated axons by the total number of axons intersected.

#### 2.5.2- Transmission electron microscopy (TEM) and image analysis

Cerebella from perfusion-fixated mice were dissected and post-fixed with 4% PFA (Sigma, P6148) and 2.5% glutaraldehyde (Sigma, G5882) in 0.1 M phosphate buffer (pH 7.4) for 1 week. Samples were then prepared at PIME platform (Cochin Institute, Paris). They were post-fixed in 2% Osmium tetroxide, dehydrated with ethanol gradient and included in epoxy resin. Semi-thin cross-section were obtained (0.5-1 µm) and stained with Methylene blue/Azur II for quality control of the appropriate region (lobule IV/V). Ultra-thin coronal slices were obtained (50-90 nm), and staining was performed on 4% aqueous uranyl acetate, flowed by Reynolds lead citrate. TEM images were acquired using a JEOL JEM-1011 microscope, and digitized with DigitalMicrograph software. Images were analyzed using ImageJ software (1.52k) after acquisition at 1200X magnification for g-ratio measurements (ratio of the inner to outer myelin sheath perimeter), and at 20 000X magnification for Major Dense Line (MDL) distance measurements, where a line was drawn from the center of one MDL to the center of the next one.

### 2.6- Behavioral tests

#### Recording material

Mice were tracked and recorded using TIS DMK 22AUC03 cameras (Stemmer) equipped with a 2.8-12 mm lens (60528, Stoelting). Any-maze software (v7.0.0) was used to record and measure mice performance.

#### Balance beam

The test was adapted from Bachsetter and colleagues (Bachstetter *et al*., 2014), and used to assess mice coordination and motor performance. Mice were separated from a housing box by a progressively narrowing beam (Campden instrument, LIC 80306): 1m in length, with the width decreasing by 10 mm every 250 mm, from an initial width of 35 mm down to 5 mm. The beam was raised to 50 cm. Mice were recorded during the crossing with cameras positioned above and beside the beam and behind the box. Test was repeated 3 times on the same day with a 10 min inter-trial-interval (ITI). The time to complete the task was measured, starting when the mouse started to move onto the bar.

#### Rotarod motor learning task

The test was adapted from Brito and colleagues (Brito *et al*., 2022). Mice were habituated to the Rotarod Orchid™ apparatus for 1 min at 4 rpm and then recorded three times a day during 5 min with 10 min ITI, and during 5 consecutive days. For the test, the speed of the rotarod was increased gradually from 4 to 40 rpm in 300 seconds (acceleration of 0.12 rpm/s). Each trial lasted until the mouse either fell from the rod or revolved 360° for three consecutive times, with a cut off of 5 min. Trials stopped when mice fell, and falls that occurring before 4 rpm were not taken into account. The latency time to fall was measured, and the average score of the three daily trials was calculated. Results represent the fall latency time of mice over five trial sessions.

#### Novel Object Recognition Test (NORT)

The NOR test aimed to assess object memory in mice as described previously (Apazoglou *et al*., 2018; Ktena *et al*., 2025). The NOR testing chamber was a transparent Plexiglass arena (openfield of 40 x 40 x 35 cm) and easily cleanable plastic objects with similar size were used. Room lighting was kept constant. One day before testing, animals underwent a habituation session for 15 min. During acquisition session, two identical objects were placed in the arena (5 cm from the wall and 10 cm apart). The 10 min-test was valid if mice spent at least 5 seconds exploring both objects. A day after, the 10 min-recognition test was conducted, allowing exploration of one familiar and one novel object. Time spent with each object and total exploration time were recorded. The Recognition Index (RI) was calculated as: (time spent with novel object x 100)/ (time spent with novel + familiar objects). A RI ≥ 55% indicates successful discrimination.

#### Social interaction test

The social interaction test was adapted from the "three chamber test" as previously described (Kaidanovich-Beilin *et al*., 2011). Mice were placed in the same arena as for the NORT, for 15 min habituation. A day after, each mouse was recorded for 10 min during free exploration of the arena containing two 10 cm-diameter mesh boxes. Active interaction, defined as sniffing a box for at least 5 seconds, was scored. In the first trial, the mouse interacted with a stranger placed under one box, while the other box was empty. In the second trial, one box contained the familiar mouse from the first trial and the other contained a novel mouse. For both trials, interaction time with each box was recorded. Climbing on the boxes without sniffing, interpreted as played or dominance behavior, was not counted as an interaction. Sociability Index (SI) was calculated as (time spent with the novel mouse x 100)/(total interaction time). A SI ≥ 55% indicates successful discrimination.

#### Y-maze

The test aims to assess the spatial working memory as previously described by Prieur and colleagues (Prieur & Jadavji, 2019). Mice were placed in an opaque Y-maze, with 120° between each arm. Arms were 8 cm-width and 39.5 cm-length. Different cues were placed on each wall for spatial differentiation. The camera above the maze tracked the mice, which were placed at the end of the arm and allowed to explore freely for 5 min. An entry was counted when all four limbs entered an arm. Spatial memory was assessed by calculating the percentage of spontaneous alternation: *i.e.* (number of correct alternations without repeat x 100)/(total arm entries-2).

### 2.7- Statistical analyses

Data were expressed as mean ± standard deviation (SD). Statistical analyses were performed using GraphPad Prism 9 software. Normality of data was assessed with the Shapiro-Wilk test and comparisons between 2 conditions were made with an unpaired Student’s t-test. For more than 2 conditions, data normal distribution was assessed using quantile plot (Q-Q plot) and two-way ANOVA followed by Sidàk’s post-hoc test was used. Rotarod data were analyzed by two-way ANOVA with repeated-measures (genotype x session) and followed by Sidàk’s post-hoc test. Differences were considered statistically for p<0.05 and indicated as the following: * for p<0.05, ** for p<0.01, *** for p<0.001, and **** for p<0.0001.

## 3- Results

### OLA10 deficiency alters OPC morphological maturation in vitro

We conducted primary cultures of OPCs to investigate the effect of OLA10 deficiency on oligodendrocyte morphology, an indicator of their maturation (Wegner, 2020). At DIV7 (**Fig. 1A**), we quantified the number of processes extending from the oligodendroglial soma and categorized cells based on their degree of ramification: low (≤ 4) or high (> 4 processes). In KO cultures, we observed a significant increase in the proportion of OPCs exhibiting low ramification compared to control (82.42% for KO vs 41.32% for control, p<0.05; **Fig. 2A**). In addition, we observed a concomitant and significant decrease in the proportion of OPCs with a high degree of ramification in KO group versus control (17.54% for KO vs 58.68% for Ctl, p<0.05; **Fig. 2A**). We also assessed the mRNA expression level of key myelin genes (*Plp*, *Mbp*, *Mag* and *Cnp*) at DIV7, and found no significant differences between KO and control groups (**Fig. 2B**).

**Figure 2:**
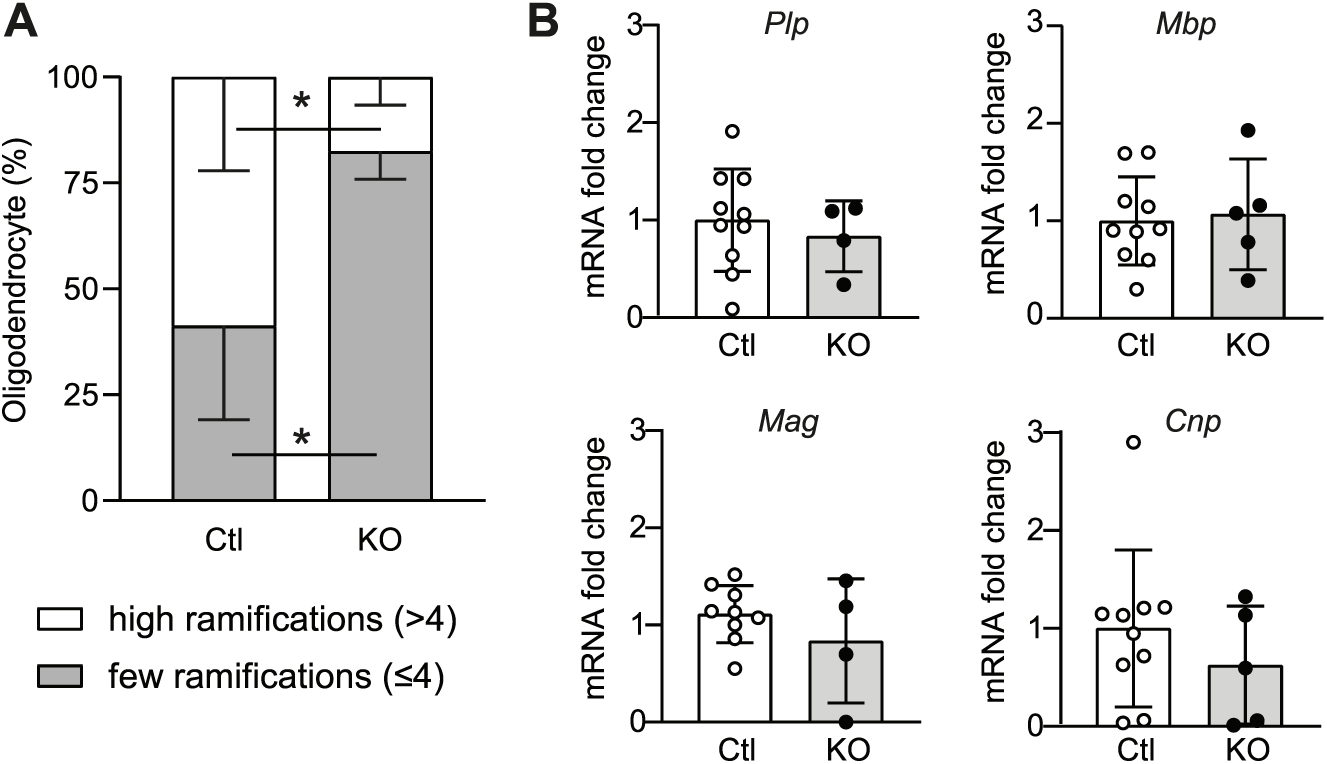
Effect of OLA10 deficiency on the morphological maturation of OPCs in primary culture. OPC primary cultures obtained from P3-4 KO and control pups were grown until DIV7. **(A)** Histogram represents the percentage of OPCs with low and high ramification in control and KO cultures at DIV7. n (Ctl/KO) = 9/3 animals from 3 independent experiments. **(B)** Histogram represents the mRNA level of myelin genes (*Plp*, *Mbp*, *Mag*, *Cnp*). n (Ctl/KO) = 8-10/4-5 animals from 3 independent experiments. Data are expressed as mean ± SD, and were analyzed using Student’s t-test. *p<0.05. *Cnp*: 2’, 3’-Cyclic-Nucleotide 3’-Phosphodiesterase; Ctl: Control; DIV: Days *in vitro; Mag*: Myelin Associated Glycoprotein; *Mbp*: Myelin Basic Protein; OPC: Oligodendrocyte Precursor Cells*, Plp*: ProteoLipid Protein; SD: Standard deviation.

### OLA10 deficiency impairs myelination in neuron/glia co-culture in vitro

To evaluate the impact of OLA10 deficiency on myelination, we performed primary neuron/glia co-cultures, and we quantified the number of axons and determined the percentage of axons that were myelinated (**Fig. 3A-C**). At DIV24 (**Fig. 1B**), the number of ADAM10^+^ oligodendrocytes were significantly decreased in KO co-cultures (**Fig. S1C-G**). We observed a significant decrease of the percentage of myelinated axons in KO group compared to controls (38% for KO vs 75% for Ctl, p<0.0001; **Fig. 3C**), while the total number of axons, neurons and ADAM10^+^ neurons remained unchanged (**Fig. 3B and Fig. S1C-E**).

**Figure 3:**
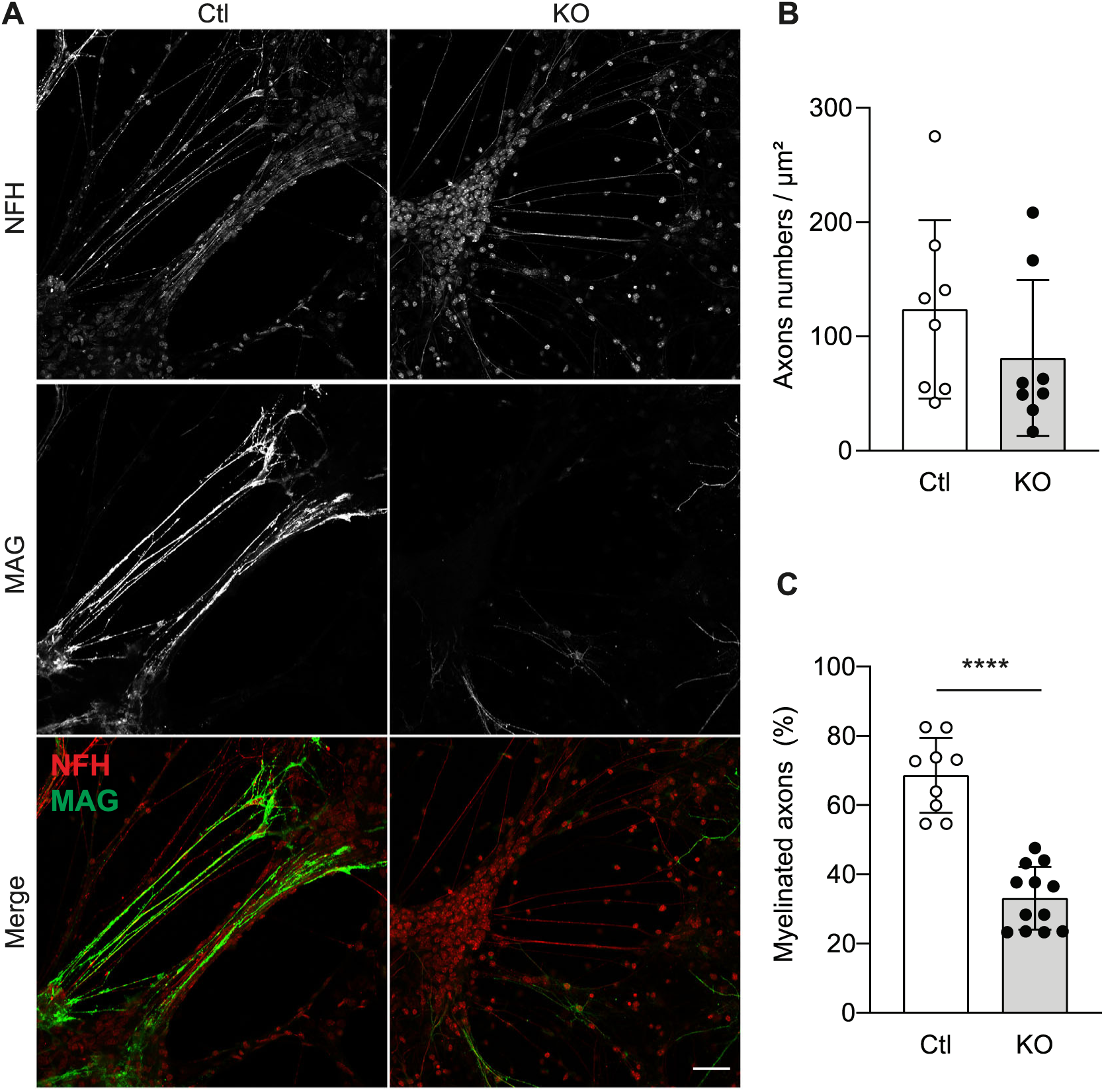
Effect of OLA10 deficiency on myelination process in primary neuron/glia co-culture. Cultures were performed with dissociated cells from E13.5 spinal cords of KOOL-A10 and control mice. Axon and myelin were labelled using Neurofilament antibody (NFH, red) and Myelin-associated glycoprotein antibody (MAG, green), respectively. **(A)** Representative confocal images of myelinated axons at DIV24. Scale bar: 20 µm. **(B, C)** Histograms represent **(B)** the number of axons per µm^2^ and **(C)** the percentage of myelinated axons. Co-labelled MAG^+^ and NFH^+^ axons were quantified to express the percentage of myelinated axons (MAG^+^NFH^+^) over total axons (NFH^+^). Data are expressed as mean ± SD and were analyzed using Student’s t-test, ****p<0.001. n (Ctl/KO) = 9/12 wells with 4/5 images/well, in 3 independent experiments. Ctl: Control; DIV: Days *in vitro;* MAG: Myelin-associated glycoprotein; NFH: Neurofilament H; SD: Standard deviation.

### OLA10 deficiency alters myelin ultrastructure in vivo

We next examined myelin ultrastructure *in vivo* to assess the effect of OLA10 deficiency on myelination 6- and 12-months following tamoxifen-induced OLA10 deficiency (**Fig. 1C**). At 6 months post OLA10 deficiency induction, we observed a significant increase in the g-ratio in the KO group compared to control group, in both male and female mice (**Fig. 4A, 4B**; male: 0.75 ± 0.08 for KO vs 0.72 ± 0.09 for Ctl; female: 0.78 ± 0.07 for KO vs 0.76 ± 0.06 for Ctl, p<0.0001) indicating that OLA10 deficiency leads to a thinner myelin sheath regardless of sex. At 12 months after OLA10 deficiency induction, the g-ratio in KO mice significantly decreased compared to their control littermates in both male and female groups (**Fig. 4C, 4D**; male: 0.70 ± 0.08 for KO *vs* 0.74 ± 0.07 for Ctl; female: 0.71 ± 0.15 for KO vs 0.738 ± 0.19 for Ctl, p<0.0001), indicating a thicker myelin sheath without any sex-related differences. To determine whether the g-ratio alterations in KO mice stemmed from changes in myelin sheath layers or modifications in the spacing between Major Dense Lines (MDL), we measured the number of sheath layers per axon as well as the distance between MDL (**Fig. 4E-G**). We observed no significant differences in the number of myelin sheath layers between male or female KO mice and their respective control group at 6 and 12 months (**Fig. 4F, 4G**). However, at 6 months after OLA10 deficiency induction, the KO groups showed a significant decrease in the distance between MDL compared to control in both sexes (**Fig. 4F**; male: 9.6 nm for KO *vs* 13.45 nm for Ctl; female: 8.5 nm for KO *vs* 10.82 nm for Ctl, p<0.0001). At 12 months, no differences in MDL distance were detected between male or female KO mice and their sex-matched controls (**Fig. 4G**).

**Figure 4:**
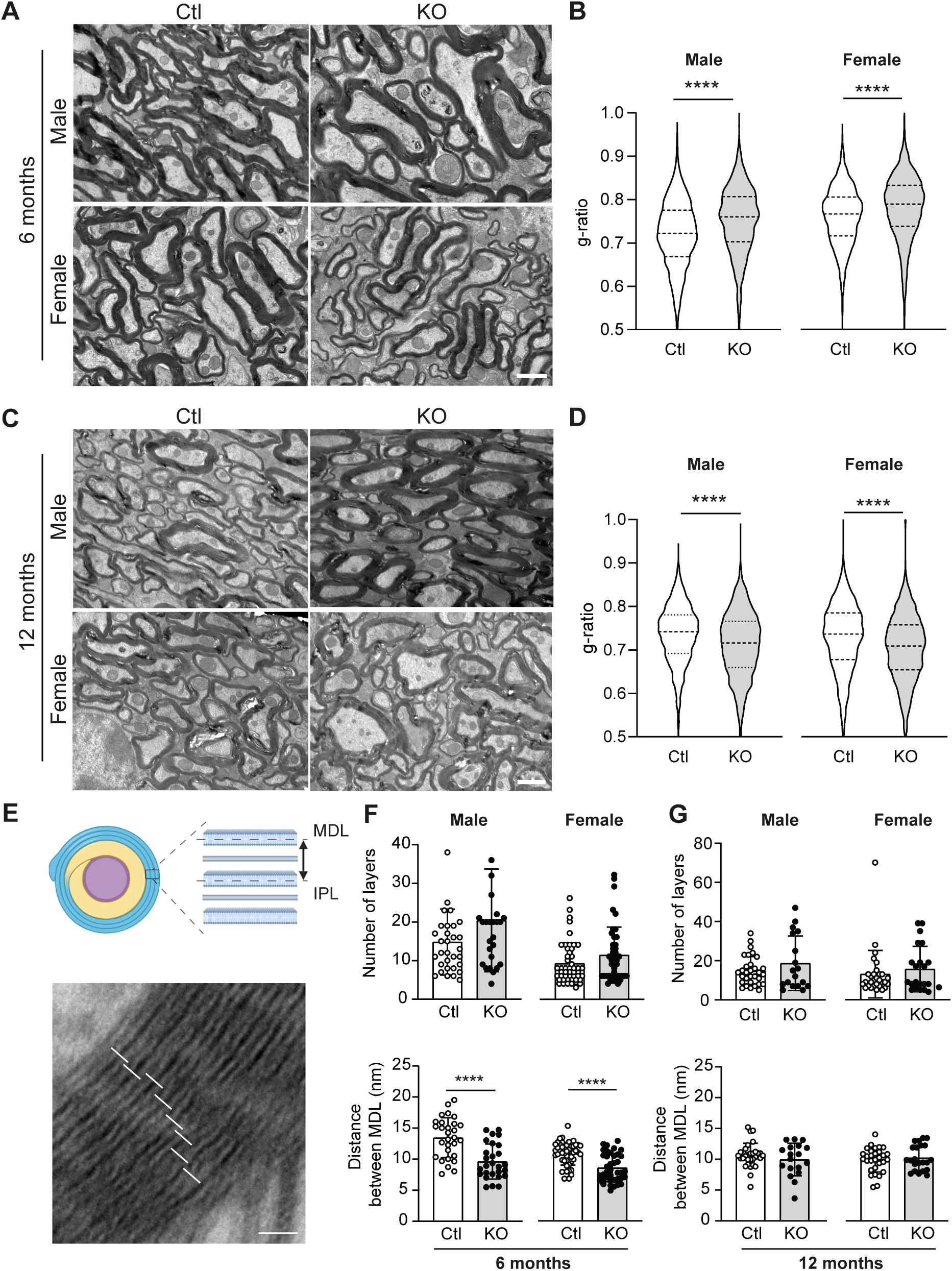
Effect of OLA10 deficiency on myelin ultrastructure *in vivo*. Adult KOOL-A10 mice were injected with tamoxifen (60 mg/kg, i.p.) during 3 consecutive days. **(A)** and **(C)** illustrate representative TEM images of transversal cross sections of cerebellum from male and female control and KOOL-A10 mice at 6 **(A)** and 12 **(C)** months after tamoxifen injection. Scale bar: 2 µm. **(B**, **D)** Histograms represent the average g-ratio of myelinated axons at 6 **(B)** and 12 **(D)** months after tamoxifen injection. **(E)** Schematic illustration of MDL distances measurement on a TEM image. Scale bar: 200 nm. **(F**, **G)** Average number of myelin sheath layers and average distance between consecutive MDLs in animals at 6 **(F)** and 12 **(G)** months following tamoxifen-induced OLA10 deficiency. Data are expressed as mean ± SD and analyzed using Student’s t-test, ****p<0.0001. (B-D) n (Ctl/KO) = 3/3 males and females with at least 1200 axons measured per group. (F) n (Ctl/KO) = 29/27 myelin sheaths in males and 48/39 in females. (G) n (Ctl/KO) = 31/18 myelin sheaths in males and 30/22 in females, with n = 3 animals/group. Ctl: Control; i.p. intraperitoneal injection, IPL: Intraperiodic Line; MDL: Major Dense Line; SD: Standard deviation, TEM: transmission electron microscopy.

### OLA10 deficiency alters MBP gene expression in vivo

To further investigate the differences observed on myelin thickness, we performed biochemical analysis to assess the effect of OLA10 deficiency on the expression of the key myelin gene MBP in several CNS tissues. Proteins and mRNAs were extracted from the brain, cerebellum and spinal cord of male and female KOOL-A10 mice, 12 months post-tamoxifen-induced OLA10 deficiency (**Fig. 1C**). We observed no significant difference in *Mbp* mRNA expression in the brain (**Fig. 5A**) whereas *Mbp* mRNA levels were significantly increased in the cerebellum (2.72-fold change for KO *vs* Ctl, p<0.05; **Fig. 5B**) and spinal cord (2.40-fold change for KO *vs* Ctl, p<0.01; **Fig. 5C**) of male KO mice. *Mbp* mRNA expression in these three regions remained unchanged in female KO mice compared to female control mice (**Fig. 5A-C**). Regarding MBP protein level, we observed a significant decrease of the 21 kDa isoform in the brain of male KO mice (70.67% for KO vs 100% for Ctl, p<0.01; **Fig. 5D, 5G**), while the 18 kDa isoform levels remained unmodified. No differences were detected in brain of female KO mice compared their control littermates. However, in the cerebellum of female KO mice, the 21 kDa isoform was significantly increased (160% for KO vs Ctl, p<0.05; **Fig. 5E, 4H**) while no effect in the 18 kDa isoform was observed (**Fig. 5K**). In the spinal cord, no significant differences in MBP protein levels were observed between KO mice and their sex-matched controls (**Fig. 5F, 5I, 5L**).

**Figure 5:**
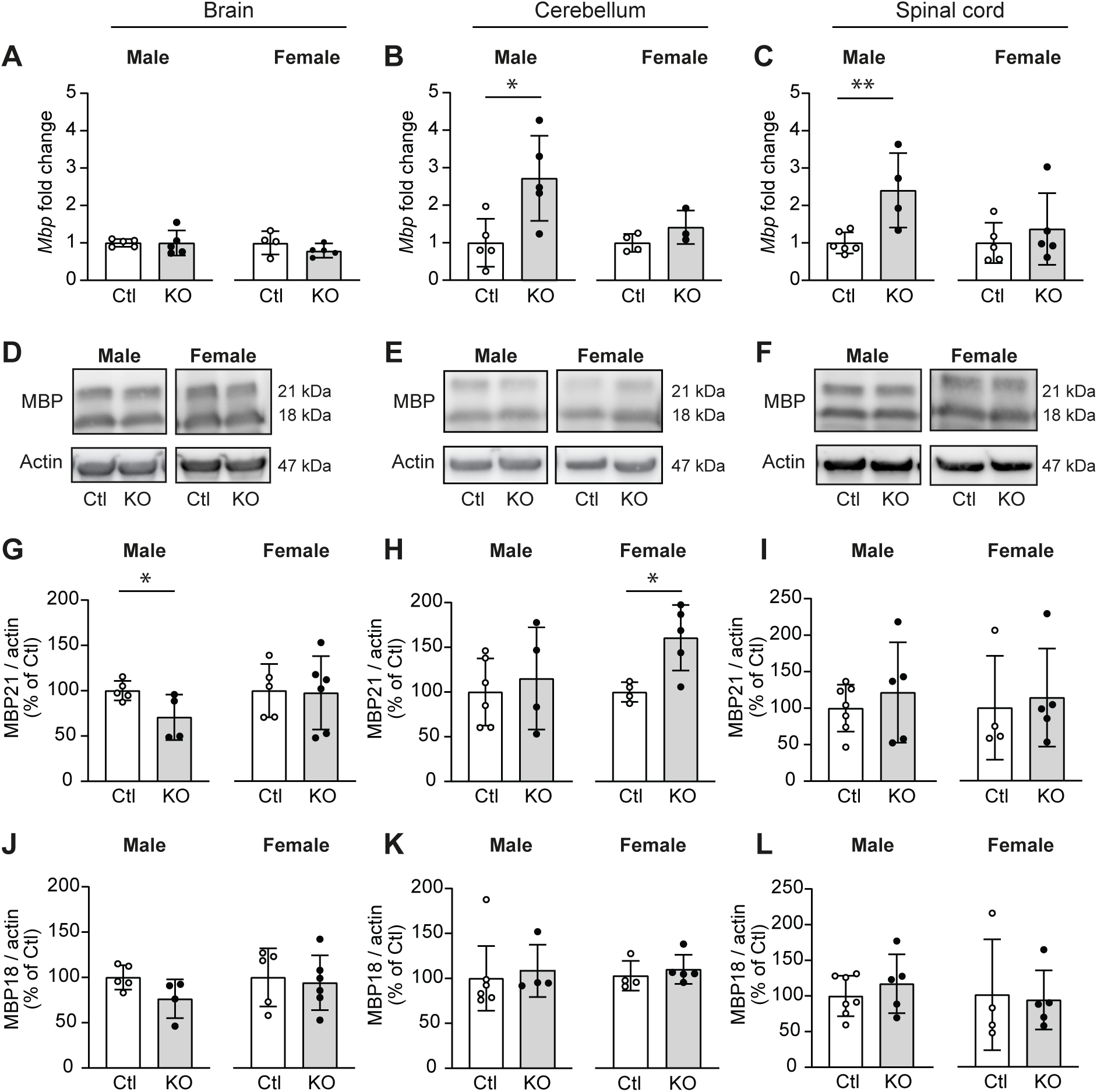
Effect of OLA10 deficiency on MBP expression. Protein and mRNA were extracted from brain, cerebellum and spinal cord of KOOL-A10 mice 12 months after tamoxifen injection. Histograms A to C indicate the expression of *Mbp* mRNA in brain **(A)**, cerebellum **(B)** and spinal cord **(C)** in male and female control and KO mice, assessed by RT-qPCR experiments. Quantification of *Mbp* mRNA was expressed in fold change. Representative western blot membranes of MBP (18 & 21 kDa isoforms) in brain **(D)**, cerebellum (**E**) and spinal cord (**F**). Actin was used as loading control. **(G-I)** Histograms of MBP (21 kDa) protein quantification in these three regions in KO and sex-matched control groups. **(J-L)** Histograms of MBP (18 kDa) protein quantification in the same groups of mice. MBP protein quantification was normalized with actin and expressed as percentage of Ctl. Data are expressed as mean ± SD. Data were analyzed using Student’s t-test. *p<0.05; **p<0.01. n (Ctl/KO) = 5-7/4-5 in males and 4-5/3-6 in females. Ctl: Control; MBP18/21: Myelin Basic Protein 18/21 kDa isoform; SD: Standard deviation.

### OLA10 deficiency impairs novel object recognition memory

To assess the impact of OLA10 deficiency on different cognitive abilities, we conducted the Novel Object Recognition test (NORT), the Social Interaction test and the Y-Maze test with spontaneous alternation (**Fig. 6A**). These tests were conducted on male and female control and KO mice, two weeks before and at 1, 6, 12 months after tamoxifen injection (**Fig. 1C**). No significant differences were observed in NORT performance up to 6 months (**Fig. 6B, 6C, 6D**). However, at 12 months, male KO showed a significant decrease in the Recognition Index (RI: 46.81% ± 11.58 for KO *vs* 66.02% ± 13.98 for Ctl, p<0.01; **Fig. 6E**), indicating impaired novel object recognition. No impairment was observed in female KO mice. Additionally, no significant differences were found between KO and control groups in either the Social Interaction test or in the Y-Maze test for both male and female (**Fig. 6F-M**).

**Figure 6:**
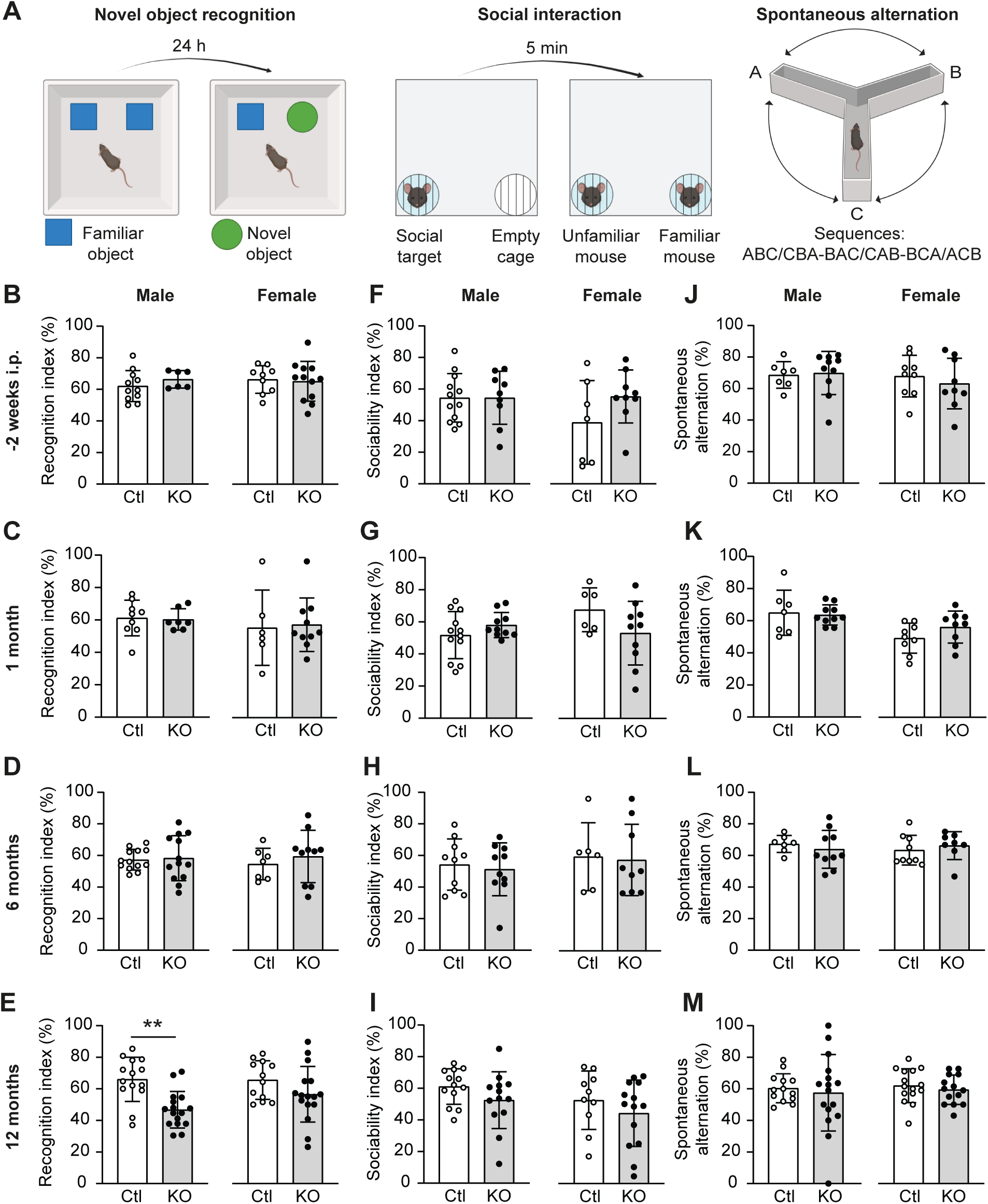
Effect of OLA10 deficiency on cognitive behavior. **(A)** Cognitive behavior tasks scheme. The NORT was performed 24 h after the habituation phase, with mice expected to discriminate a novel object from a familiar one. In the Social interaction test, mice were expected to discriminate between a familiar and a non-familiar mouse. In Y-maze test, mice were expected to spontaneous alternate between arms, avoiding the one most recently visited. **(B-E)** Histograms represent performance in the NORT, expressed as a Recognition Index percentage, two weeks prior tamoxifen **(B)**, and at 1 **(C)**, 6 **(D)**, 12 **(E)** months after tamoxifen injection in KO and sex-matched control groups. **(F-I)** Histograms represent the percentage of Sociability Index, two weeks prior tamoxifen **(F)**, and at 1 **(G)**, 6 **(H)**, 12 **(I)** months after tamoxifen injection in the same groups. **(J-M)** Histograms represent the spatial memory capacity as a percentage of spontaneous alternation 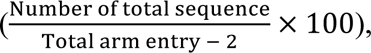 two weeks prior tamoxifen **(J)** and at 1 **(K)**, 6 **(L)**, 12 **(M)** months after tamoxifen injection in the same groups of mice. Data are expressed as mean ± SD. Data were analyzed using Student’s t-test, **p<0.01. (B-D) n (Ctl/KO) = 10/6 males and 9/12 females, (E) n (Ctl/KO) = 14/15 males and 11/16 females. (F-H) n (Ctl/KO) = 12/9 male and 7/9 females, (I) n (Ctl/KO) = 13/13 males and 10/14 females. (J-L) n (Ctl/KO) = 7/10 males and 9/9 females, (M) n (Ctl/KO) = 13/15 males and 14/14 females. Ctl: Control; i.p.: intraperitoneal injection of tamoxifen; NORT: Novel Object Recognition Test; SD: Standard deviation.

### OLA10 deficiency alters fine motor activity

The balance beam test was used to assess fine motor coordination in KOOL-A10 mice (**Fig. 7A**). Animals were required to cross a progressively narrowing beam two weeks before, and at 1, 6 and 12 months following tamoxifen-induced OLA10 deficiency (**Fig. 1C**). Prior to invalidation, no differences were observed between KO mice and their matched controls (**Fig. 7B**). From 1 to 12 months post tamoxifen, male KO mice exhibited a significant increase in the time taken to cross the beam compared to control littermates (1 month: 30.84s ± 10.66 for KO *vs* 14.26s ± 6.29 for Ctl, p<0.001; 6 months: 33.81s ± 7.68 for KO *vs* 24.39 ±6.56s for Ctl, p<0.01; 12 months: 44.28s ± 8.91 for KO *vs* 29.23s ± 12.09 for Ctl, p<0.05). In contrast, no significant differences were observed in female KO mice compared to their controls (**Fig. 7C-E)**.

**Figure 7:**
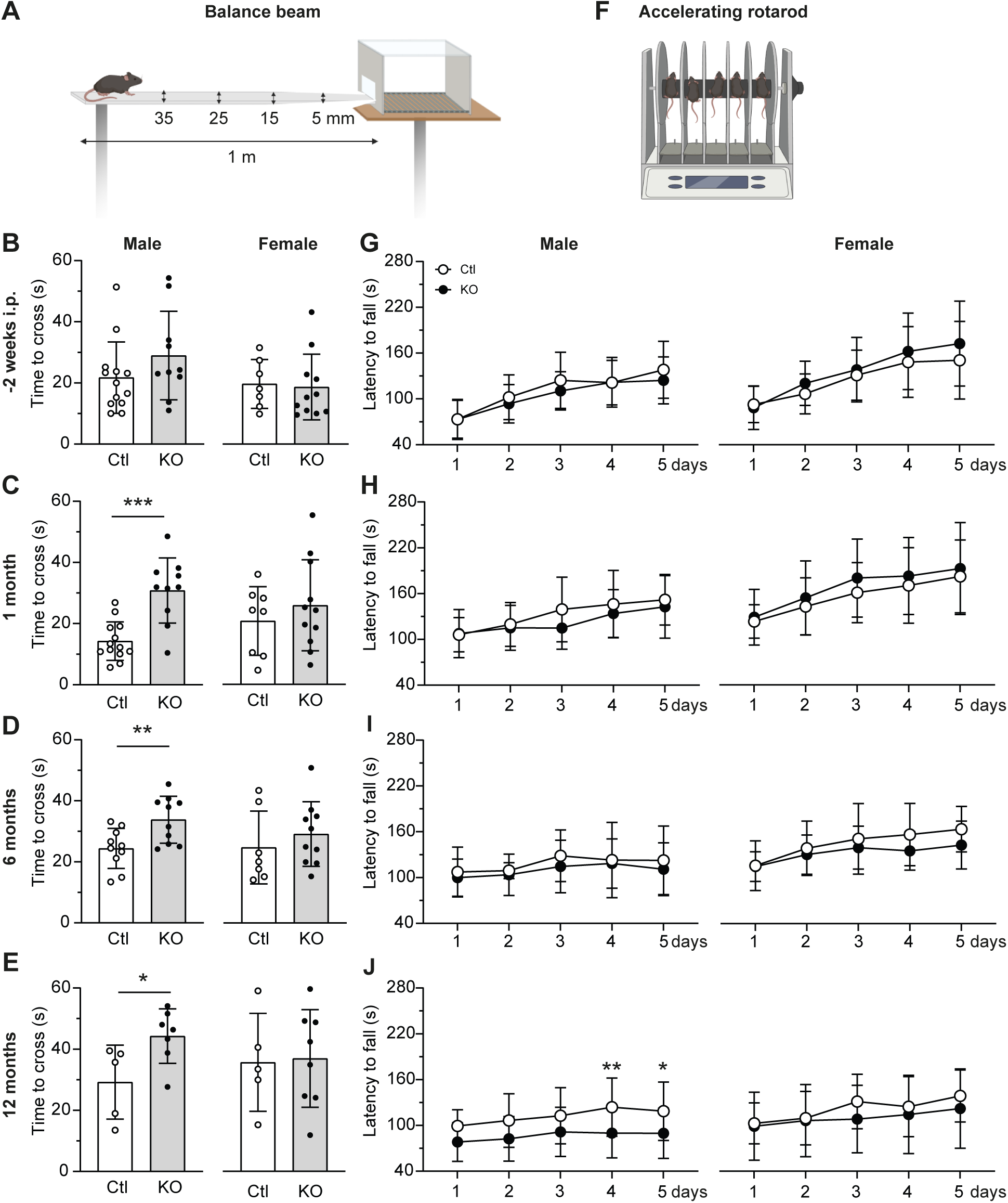
Effect of OLA10 deficiency on motor behavior. **(A)** The balance beam is a fine motor test in which mice were expected to cross a narrowing bar to reach a safe zone. The accelerating rotarod is a motor learning task in which mice were expected to remain on an accelerating rod over the 5 consecutive days the test is performed. **(B-E)** Histograms of balance beam performance in KO and sex-matched control groups. Time to cross the bar was used to assess performance prior tamoxifen injection **(B)**, and at 1 **(C)**, 6 **(D)** and 12 **(E)** months after. **(F)** Assessment of performances in accelerating rotarod in the same groups of mice. Latency to fall from the rod was used to assess motor learning abilities. Performances were evaluated daily during the 5 day-test period before tamoxifen injection **(G)** and at 1 **(H)**, 6 **(I)** and 12 **(J)** months after. Data were presented as mean ± SD and were analyzed using Student’s t-test (B-E) and two-way ANOVA with Sidàk post-hoc test (G-J). (B-D) n (Ctl/KO) = 12/10 males and 7/11 females, (E) n (Ctl/KO) = 5/7 males and 5/8 females, (F) and (H) n (Ctl/KO) = 17/19 males and 18/19 females, (G) n (Ctl/KO) = 17/19 males and 18/19 females, (I) n (Ctl/KO) = 7/10 males and 9/9 females, (J) n (Ctl/KO) = 17/21 males and 12/14 females, *p<0.05, **p<0.01, ***p<0.001. Ctl: Control, i.p.: intraperitoneal injection of tamoxifen, SD: Standard deviation.

### OLA10 deficiency alters motor learning

We performed the repeated accelerating rotarod test to assess coordination and learning motor abilities in KOOL-A10 mice (**Fig. 7C**). Mice were required to maintain their position on an accelerating rotating rod, and the test was conducted over 5 consecutive days in KO and sex-matched control groups. The test was performed two weeks before and at 1, 6, 12 months after induction of OLA10 deficiency (**Fig. 1C**). No differences were observed between KO and control groups prior tamoxifen, and at 1- and 6-months post-invalidation (**Fig. 7G-I**). However, at 12 months after OLA10 deficiency induction, male KO mice showed significant impairments after 4 days (4^th^ day: 89.82s ± 32.31 for KO *vs* 123.84s ± 38.17 for Ctl; p<0.01) and 5 days (5^th^: 89.69s ± 32.76 for KO *vs* 118.61s ± 38.35 for Ctl; p<0.05), whereas no differences were observed in female groups (**Fig. 7J**).

## 4- Discussion

We had previously shown the involvement of α-secretases in neuroprotection (Siopi *et al*., 2013), in oligodendrocyte maturation processes and myelin repair after CNS injury (Llufriu-Dabén *et al*., 2018, 2019; Carrete *et al*., 2021) and have recently described the mapping of ADAM10 protein expression in the CNS highlighting ADAM10 expression in oligodendrocytes besides neurons (Padilla-Ferrer *et al*., 2024). The present study assessed the role of OLA10 in myelination and myelin maintenance using a mouse line that we generated, KOOL-A10, where OLA10 deficiency can be induced upon tamoxifen treatment at the desired time point. Our longitudinal study pointed out behavioral deficits and myelin ultrastructural changes, as well as impairments in oligodendrocyte development and myelination *in vitro* after OLA10 deficiency.

We studied the process of oligodendrocyte development *in vitro* based on the analysis of oligodendrocyte branching which reflects their maturation stage. In fact, the process of differentiation involves morphological transformation characterized by the development of a network of branches, accompanied by the expression of specific cellular markers and transcription factors (Wegner, 2020). We observed a reduced number of highly branched oligodendrocytes in KO primary culture of OPCs that reveals the impact of ADAM10 deficiency on their maturation. Differentiation of OPCs into more mature oligodendrocytes would therefore be slowed down by ADAM10 deficiency. These data are in agreement with our previous results showing that activation of α-secretases by etazolate increases OPC branching in mixed glial primary culture and the effect is reversed in the presence of a potent ADAM10 inhibitor (Llufriu-Dabén *et al*., 2018). The present results obtained on OLA10-deficient OPCs therefore confirm and extend our previous pharmacological data, revealing that oligodendroglial ADAM10 is the α-secretase that plays a key role on the maturation of oligodendrocytes.

We performed molecular analysis of *Plp*, *Mbp*, *Mag* and *Cnp* mRNAs levels in OLA10-deficient cells *in vitro* and observed no change. Therefore, OLA10 deficiency does not impact the expression of myelin genes under our experimental conditions. This lack of variation, together with the observed morphological alterations, suggests that OLA10 loss may alternatively, lead to an impairment in cell morphology and processes outgrowth, through cytoskeletal dysfunction. Indeed, evidence suggests that ADAM10 regulates cytoskeleton dynamics in various cells including neural precursor cells and neurons, influencing cell migration, morphology and neurite outgrowth (Klingener *et al*., 2014; Dreymueller *et al*., 2017; Sanz *et al*., 2017; Yang *et al*., 2017; Hsia *et al*., 2021). These effects may be mediated either directly through the cleavage of adhesion proteins or extra-cellular matrix components (Klingener *et al*., 2014; Rosenbaum & Saftig, 2024) or indirectly, for example, through the ADAM10 release of sAPPα (Chasseigneaux *et al*., 2011; Hida *et al*., 2025) or the Notch intracellular domain (NICD), which can subsequently regulate microtubule stability in cortical neurons (Yang *et al*., 2017) or can be translocated to the nucleus to modulate target genes involved in OPC differentiation, such as Hes1, 5 or Id2, 4 (Brosnan & John, 2008; Juryńczyk & Selmaj, 2010; Li *et al*., 2021). Interestingly, a role for ADAM10 in OPC maturation has been suggested in zebrafish, notably via the cleavage of the ADAM10 substrate protein L1 (Linneberg *et al*., 2019). Our present data therefore confirm the hypothesis of a pro-differentiating role for ADAM10 in OPCs.

We then aimed to determine whether this altered OPC maturation has an impact on myelination. Our results obtained on neuron/glia co-cultures revealed a reduction in the number of myelinated axons in KO cultures. This reduction was not due to a decreased number of oligodendrocytes or neurons (**Fig. S1**), and could therefore be attributed to OLA10 deficiency. OLA10 thus appears to play a positive role on myelination capacity of oligodendrocytes. It should be noted that a recent study using NG2-promotor-driven Cre recombinase showed that, *in vivo*, the ADAM10 deficiency first leads to an early myelination, and subsequently an impairment in myelin maintenance (Guo *et al*., 2022). However, this mouse model differs from ours. First, these mice have a limited life expectancy: they die at around P65 and, second, NG2-promotor-driven OLA10 deficiency is constitutive and not inducible as ours, preventing time-controlled analysis of deletion effect. Additionally, since NG2 is also expressed in neurons, astrocytes and pericytes in addition to oligodendrocytes (Viganò & Dimou, 2016), this model could not be entirely cell specific, and the observed effects could partially result of ADAM10 deletion in different brain cell types. Our model has the advantage to be more oligodendrocyte specific since we used the PLP promotor to generate our KOOL-A10 mouse model. In fact, the PLP promotor is active in OPCs and particularly in mature oligodendrocytes (Tuason *et al*., 2008). Meanwhile, another study further supports the involvement of ADAM10 in myelin sheath development, but in the particular context of cuprizone-induced demyelination (Zhu et al., 2021). The authors report that a non-cell-specific overexpression of ADAM10 via AAV injection into the corpus callosum enhances *in vivo* remyelination. Furthermore, they show that overproduction of sAPPα, the product of APP cleavage by α-secretases, induces similar effects. We have also previously shown that sAPPα treatment of cerebellar slice cultures prevents lysolecithin-induced demyelination *ex vivo* (Llufriu-Dabén *et al*., 2018). Interestingly, recent study demonstrated that oligodendrocytes can produce sAPPα via ADAM10 (Sasmita *et al*., 2024; Hida *et al*., 2025). APP seems essential for the proper myelin formation *in vivo*, as its absence shortens the distance between nodes of Ranvier (Xu *et al*., 2014), and leads to significant hypomyelination and thinner myelin sheaths (Truong *et al*., 2019). Therefore, it is plausible that the effects observed in the absence of OLA10 result from the lack of APP-derived sAPPα. The impairment in myelination observed upon OLA10 deficiency *in vitro* prompted us to examine myelin ultrastructure following the induction of OLA10 deficiency in adult mice *in vivo*. We examined cerebellar myelin ultrastructure 6 and 12 months after tamoxifen injection. At 6 months, KO animals exhibited an increased g-ratio alongside a reduced distance between MDLs, indicating thinner myelin sheaths possibly resulting from enhanced compaction. Interestingly, at 12 months, the g-ratio decreased in KO animals, suggesting a thicker myelin sheath or possible decompaction, although no significant changes were detected in the number of layers or the distance between MDLs. Thinner myelin sheaths at 6 months may indicate hypomyelination, or transient demyelination followed by incomplete remyelination. This pattern has been well documented in animal models like the cuprizone model, where early remyelination produces myelin sheaths thinner than the original (Duncan *et al*., 2017). Late thickening at 12 months may potentially reflect a compensatory response, with an excess production of myelin to stabilize neural circuits weakened by the initial phase of deficient myelination. Interestingly, myelin sheath compaction is mediated in part by the MBP protein within the oligodendrocyte plasma membrane (Vassall *et al*., 2016; Raasakka *et al*., 2017; Träger *et al*., 2020), and notably, MBP can be cleaved by ADAM10 (Amour *et al*., 2000). Remarkably, invalidation of ADAM17 in Schwann cells has been shown to alter myelin sheath thickness and reduce MBP production (La Marca *et al*., 2011).

As previously said, since APP is also a substrate of ADAM10 and its KO leads to altered myelin sheath thickness and hypomyelination in both the CNS and PNS, these findings collectively support the hypothesis of an interconnection between ADAM10, its main substrate APP, and myelin compaction. Moreover, the fact that OLA10 deficiency is induced in adulthood, when myelin is stabilized, and still causes myelin alteration, reveals a role for OLA10 in the maintenance of myelin.

To better understand the myelin impairment after OLA10 deficiency *in vivo*, we analyzed MBP expression across three CNS regions. MBP exists in several isoforms (21, 18, 17.5 and 14 kDa) (Raasakka & Kursula, 2020). The 18.5 kDa isoform plays a key role in myelin compaction, while the 21.5 kDa isoform can translocate to the nucleus and may contribute to oligodendrocyte differentiation and proliferation (Harauz & Boggs, 2013; Smith *et al*., 2013; Ozgen *et al*., 2014). Twelve months after OLA10 invalidation, a decrease in the 21 kDa MBP isoform was observed in the male brain, whereas *Mbp* mRNA levels were increased in the cerebellum and spinal cord of KO males. In contrast, only an increase in the 21 kDa MBP isoform was detected in the cerebellum of KO females.

The last part of our study focused on the behavioral (motor and cognitive) consequences of OLA10 deficiency over time. We assessed the cognitive behavior of KOOL-A10 mice using different tests in male and female mice. While no impairment was found in the Y-Maze and Social interaction tests, we cannot exclude subtle behavioral changes in social behavior or spontaneous spatial memory that might have been detected using additional tests.

However, a decline in novel object recognition memory was observed in KO male, 12 months after OLA10 deficiency. Although oligodendrocytes and/or myelin alterations are usually associated with motor dysfunction, our findings indicate that OLA10 deficiency can also result in recognition memory impairments, highlighting cognitive consequences that extend beyond motor deficits. Another study showed that OPC development is required for non-motor learning and cognition in working memory tasks (Shimizu et al., 2023). Additionally, some reports have linked myelin alterations to long-term memory-deficits, in demyelinating context or diseases (Cho et al., 2023; Mercier et al., 2024). Our findings align with, and further support, growing evidence indicating that oligodendrocytes contribute to multiple learning and memory-related processes (Munyeshyaka & Fields, 2022). Finally, we also observed alterations in fine motor behavior in KO males as early as one month after invalidation and persisting up to 12 months, as well as impaired motor learning at 12 months. These deficits could be attributed to the myelin compaction changes observed at 6 and 12 months, as these structural myelin alterations are known to induce motor disorders (Bloom et al., 2022). The fact that our model affects motor behavior earlier than learning ability may reflect a differential sensitivity of the neural circuits involved. Overall, our data point to a role for ADAM10 not only in motor functions, but also in memory, both of which depending on the integrity of oligodendrocytes and myelin (McKenzie et al., 2014; Pan et al., 2020; Steadman et al., 2020; Alberini, 2025; Angulo, 2025). Our data highlight deficits that can be induced in adulthood and appear over the long term, suggesting compromised myelin maintenance and indicating that certain phenotypes emerge only at later stages. Supporting this, several studies demonstrated that continuous generation of oligodendrocytes and myelin renewal throughout life is crucial for white matter maintenance and plasticity (Tripathi et al., 2017; Meschkat et al., 2022).

Importantly, as our model does exhibit OLA10 deficiency, with approximately a 50% decrease in its expression (**Figs. S1, S2**), it also indicates that the invalidation is not total, a result that could be expected as Cre/Lox recombination activity rarely achieves complete gene deletion (Feil et al., 2009; Jahn et al., 2018), with some cells potentially escaping Cre recombinase activity (Song & Palmiter, 2019). Nevertheless, despite this partial invalidation, we observe clear cellular, histological and functional alterations, demonstrating that this deficiency is sufficient to elicit biologically significant deficits.

A particularly interesting aspect of our results is also the evidence of sex differences, with effects observed specifically in males. Previous studies have documented sex-related differences in myelin content, structure, rate turnover and response to injury. For example, in the corpus callosum, males exhibit more myelinated axons, with thicker sheaths and shorter internodes compared to females, differences largely attributed to androgen influence (Cerghet et al., 2009; Abi Ghanem et al., 2017; Chowen & Garcia-Segura, 2021). Moreover, females exhibit higher rates of both oligodendrocyte generation and apoptosis compared to males, suggesting that myelin is more dynamically replaced in females (Cerghet *et al*., 2009). As highlighted by recent literature (Paus, 2009; Seeker & Williams, 2022; Autio-Kimura et al., 2024; Khodanovich et al., 2024; Nesbitt et al., 2024; Poulen et al., 2025), the sexual dimorphism we highlighted is of particular importance in the current research context and warrants further investigation in future studies.

In conclusion, our results indicate that OLA10 deficiency leads to defects in both oligodendrocyte maturation and myelin formation, as well as impairments in myelin maintenance. We hypothesize that oligodendrocytes exhibiting delayed morphological maturation and reduced myelination capacity in vitro, contributing in turn to myelin sheath alteration, which may have long-term consequences on myelin maintenance and lead to behavioral deficits. Our study thus highlights the subtle yet significant role of oligodendrocyte-expressed ADAM10 in regulating myelin maintenance and dynamics over time. As emerging evidence points to a role for ADAM10 in myelination and remyelination processes (Klingener et al., 2014; Zhu et al., 2021; Guo et al., 2022), our findings not only support this view but also extend it by demonstrating that ADAM10 expressed specifically in oligodendrocytes could be a key driver of these processes.

## Supporting information

Supplemental Lavaud 2025

## Abbreviations

ADAM: A Disintegrin And Metalloproteinase
APP: Amyloid Precursor Protein
CaBP: Calbindin Protein
CNP: 2’3’-Cyclic Nucleotide 3’-Phosphodiesterase
CNS: Central Nervous System
Ctl: Control
DIV: Days *in vitro*
E: Embryonic day
FGF: Fibroblast Growth Factor
GAPDH: Glyceraldehyde-3-Phosphate Dehydrogenase
Hsp60: Heat shock protein 60
i.p.: intraperitoneal injection
ITI: Inter-Trial-Interval
KOOL-A10: oligodendroglial knock-out for ADAM10
MAG: Myelin-Associated Protein
MDL: Major Dense Line
MBP: Myelin Basic Protein
Mrlp10: Mitochondrial Ribosomal Protein L10
NG2: Neuronal Glial antigen 2
NICD: Notch Intracellular Domain
OLA10: Oligodendroglial ADAM10
OPC: Oligodendrocyte Precursor Cell
P: Postnatal day
PBS-GTA: Phosphate Buffer Saline with Gelatin, Triton Azide
PDGFRα: Platelet-Derived Growth Factor receptor alpha
PLP: Proteolipid Protein
PPIA: Peptidylprolyl isomerase A
RT: Room temperature
RT-qPCR: Reverse Transcription-quantitative Polymerase Chain Reaction
SD: Standard deviation

## Acknowledgments

We acknowledge the non-profit organizations “Fondation des Gueules Cassées” (FGC), ARSEP foundation (Association pour la Recherche sur la Sclérose En Plaques), Université Paris Cité and Inserm (Institut national de la santé et de la recherche médicale) for financially supporting this work. We acknowledge the core facility of BioMedTech Facilities INSERM US36 | CNRS UAR2009 | Université Paris Cité for Imaging facility (F. Licata), Biochemical facility (C. Tomkiewicz), Cyto2BM (S. Dupuy) and Animal facility (I. Blanchard, L. Duhamel and B. Coqueran). We are very grateful to D. R. Ulusoy and G. Ríos Concepción for their help with cell culture and branching quantification. We also thank the PIME facility at Cochin Institute (Dr. A. Schmidt). APF (2019-22) and PA (2023-26) received a MENRT fellowship and ML was a recipient of an FGC fellowship (2022-25).

## Notes

### Competing Interest Statement

The authors have declared no competing interest.

