## Supplemental Lavaud 2025 for "Role of oligodendroglial ADAM10 in oligodendrocyte maturation, myelination and myelin maintenance"

### Supplemental information

**Figure S1: Tamoxifen-induced ADAM10 deficiency in oligodendrocytes *in vitro***
OPCs were isolated from P3-P4 KOOL-A10 mice treated at P0 with tamoxifen (100 mg/kg), and cultured until DIV7 with 4-OH-tamoxifen (1 µM). **(A)** *Adam10* expression was assessed by quantifying mRNA with three set of primers designed upstream (exons 1-2) or downstream (exons 6-7 or 13-14) the excised exon upon tamoxifen treatment. **(B)** ADAM10 protein expression was quantified using western-blot. Two ADAM10 bands were detected, the precursor form (85 kDa) and the mature form (65 kDa). Actin was used as a loading control. **(C)** Representative confocal images from DIV24 primary neuron/glia co-culture using E13.5 KOOL-A10 embryos. Immunostaining was performed for ADAM10 (red), oligodendrocyte (Olig2, green) and neuron (NeuN, blue). Immunolabelled $Olig2^+$ cells were indicated by full arrowheads, ADAM10 and Olig2 co-labelled cells ($A10^+Olig2^+$) by empty arrowheads, and ADAM10 and NeuN co-labelled cells ($A10^+NeuN^+$) by full arrows. Scale bar: 20 µm. Quantitative analysis of **(D)** the number of $NeuN^+$ cells per $\mu m^2$; and the percentage of **(E)** $ADAM10^+$ $NeuN^+$ cells over $NeuN^+$ cells, **(F)** the number of $Olig2^+$ cells per $\mu m^2$ and the percentage of **(G)** $ADAM10^+$ $Olig2^+$ cells over $Olig2^+$ cells. Data were analyzed using Student's t-test. (A) n (Ctl/KO) = 8-7/5 animals from 3 independent experiments; (B) n (Ctl/KO) = 7/15 animals from 3 independent experiments; (C-G) n (Ctl/KO) = 8/8 culture wells with at least 3 pictures/well in 3 independent experiments; *$p<0.05$; **$p<0.01$. A10: ADAM10; Ctl: Control, DIV: Days *in vitro*; NeuN: Neuronal Nuclear Antigen; Olig2: Oligodendrocyte Transcription Factor 2; SD: Standard deviation.

**Figure S2: Tamoxifen-induced ADAM10 deficiency in oligodendrocytes *in vivo***
**(A)** *Adam10* mRNA quantification in adult brain KOOL-A10 mice. Histograms represent *Adam10* mRNA fold change in males and females control and KO groups (exons 6-7). Male and female KOOL-A10 adult mice were treated with tamoxifen (60 mg/kg; i.p.) during 3 consecutive days and brains were collected 12 months post tamoxifen injection and subjected to RT-qPCR. n (Ctl/KO) n=5-6/5 males and n (Ctl/KO) =4-5/4-6 females. **(B)** ADAM10 protein quantification assessed by western-blot. Proteins were extracted from brain mice at 12 months post tamoxifen injection. n (Ctl/KO) = 6/5 males and 5/6 females. **(C)** Representative confocal images of thalamic regions from brain coronal sections of P21 KOOL-A10 mice treated with tamoxifen at P0 (100 mg/kg). Immunostaining was performed for ADAM10 (red), oligodendrocytes (Olig2, green) and neurons (NeuN, blue). Immunolabelled $Olig2^+$ cells were indicated by full arrowheads, ADAM10 and Olig2 co-labelled cells ($A10^+Olig2^+$) by empty arrowheads, and ADAM10 and NeuN co-labelled cells ($A10^+NeuN^+$) by full arrows. Scale bar: 20 µm. Quantitative analysis of **(D)** the number of $Olig2^+$ cells per $\mu m^2$; the percentage of **(E)** $ADAM10^+$ $Olig2^+$ cells over $Olig2^+$ cells and **(F)** $ADAM10^+NeuN^+$ cells over $NeuN^+$ cells. Data are expressed as mean ± SD and analyzed using Student's t-test. *$p<0.05$; **$p<0.01$. Quantification of *Adam10* mRNA was expressed in fold change and ADAM10 protein quantification was normalized to actin and expressed as percentage of Ctl. (C-F) n (Ctl/KO) = 4/4 males and females with at least 4 images/animal. A10: ADAM10; DIV: Days *in vitro*, NeuN: Neuronal Nuclear Antigen; Olig2: Oligodendrocyte Transcription Factor 2; SD: Standard deviation.

*Figure S1*

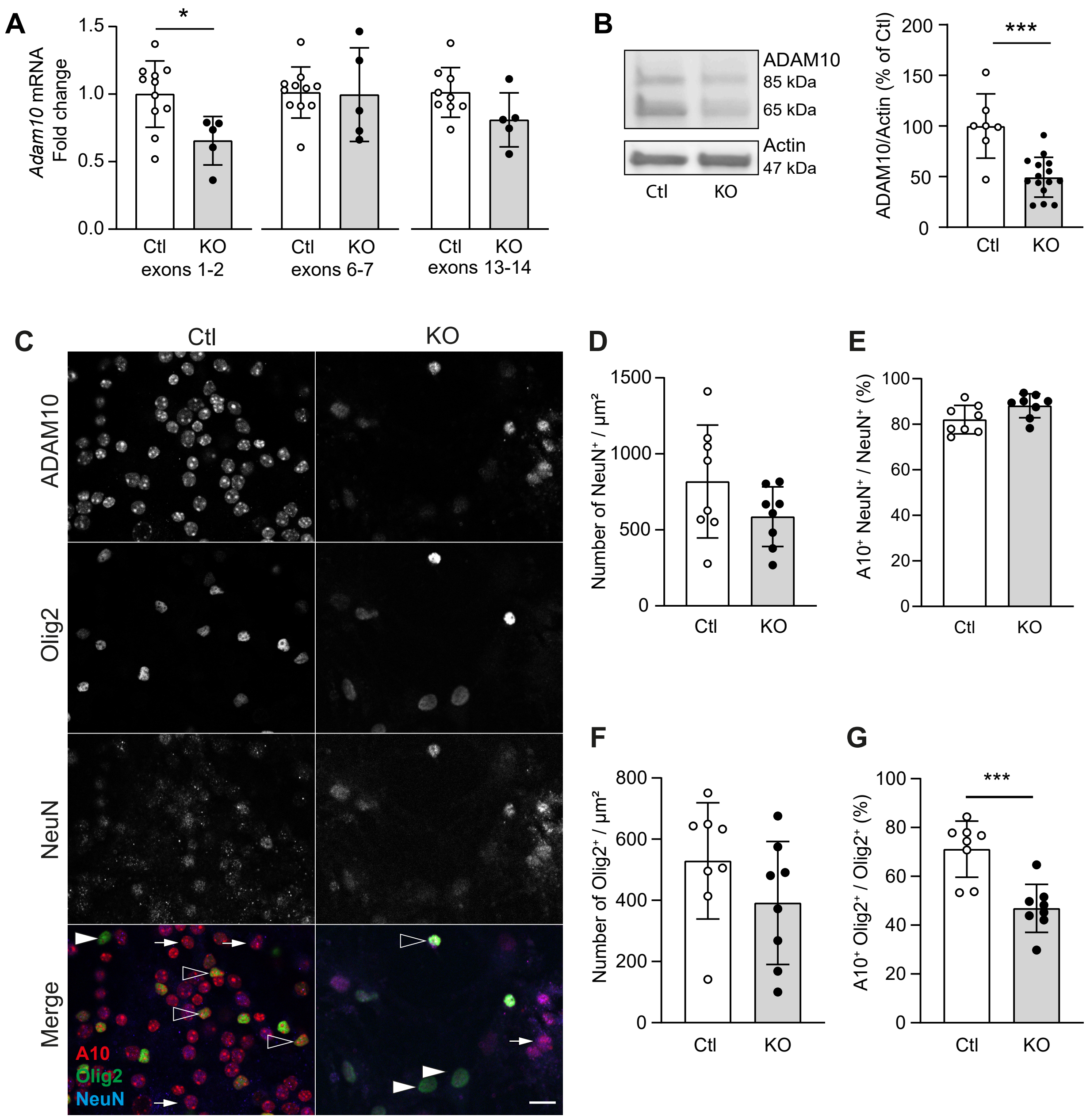
A
Adam10 mRNA
Fold change
1.5
1.0
0.5
0.0
*
Ctl
KO
exons 1-2
Ctl
KO
exons 6-7
Ctl
KO
exons 13-14
B
ADAM10
85 kDa
65 kDa
Actin
47 kDa
Ctl
KO
ADAM10/Actin (% of Ctl)
200
150
100
50
0
***
Ctl
KO
C
Ctl
KO
ADAM10
Olig2
NeuN
Merge
A10
Olig2
NeuN
D
Number of NeuN⁺ / µm²
1500
1000
500
0
Ctl
KO
E
A10⁺ NeuN⁺ / NeuN⁺ (%)
100
80
60
40
20
0
Ctl
KO
F
Number of Olig2⁺ / µm²
800
600
400
200
0
Ctl
KO
G
A10⁺ Olig2⁺ / Olig2⁺ (%)
100
80
60
40
20
0
***
Ctl
KO

*Figure S2*

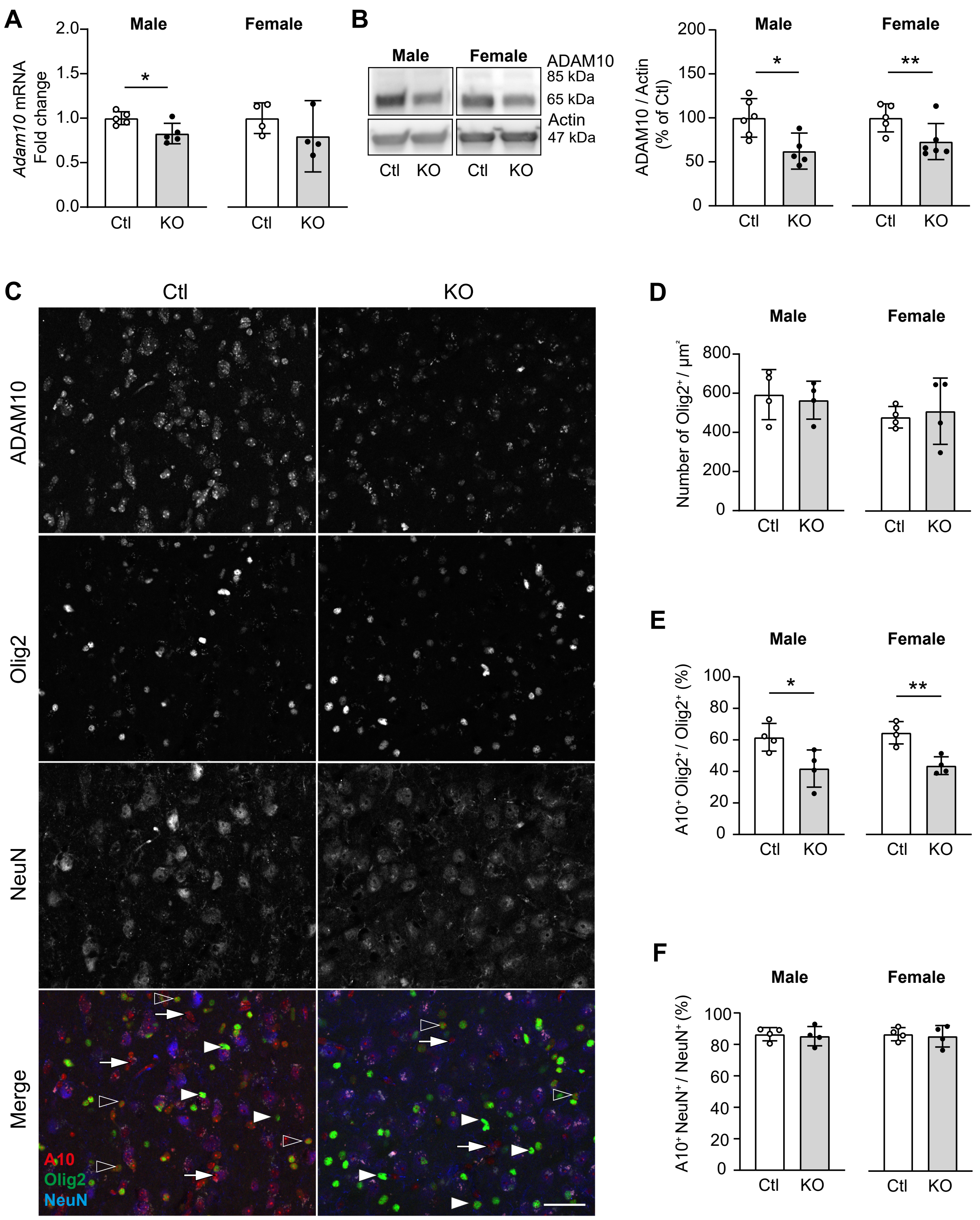

A
Male
Female
Adam10 mRNA Fold change
2.0
1.5
1.0
0.5
0.0
*
Ctl
KO
Ctl
KO
B
Male
Female
ADAM10
85 kDa
65 kDa
Actin
47 kDa
Ctl
KO
Ctl
KO
ADAM10 / Actin (% of Ctl)
200
150
100
50
0
*
**
C
Ctl
KO
ADAM10
Olig2
NeuN
Merge
A10
Olig2
NeuN
D
Male
Female
Number of Olig2+ / µm²
800
600
400
200
0
E
Male
Female
A10+ Olig2+ / Olig2+ (%)
100
80
60
40
20
0
*
**
F
Male
Female
A10+ NeuN+ / NeuN+ (%)
100
80
60
40
20
0
